# Crb3 and NF2: A dynamic duo that controls assembly of the apical junctions and barrier function via Rho/ROCK signaling

**DOI:** 10.1101/2025.01.15.633166

**Authors:** S. Fan, S. Varadarajan, V. Garcia-Hernandez, B. Margolis, C.A. Parkos, A. Nusrat

**Affiliations:** Department of Pathology, University of Michigan Medical School, Ann Arbor, MI 48109, USA; Department of Internal Medicine, University of Michigan Medical School, Ann Arbor, MI 48109, USA

**Author notes:** Corresponding authors: Correspondence:Asma Nusrat, University of Michigan Medical School Department of Pathology, 4057 BSRB, 109 Zina Pitcher Place, Ann Arbor, MI 48109, USA., Charles A Parkos, University of Michigan Medical School Department of Pathology, NCRC, Building 35, Rm 30-1537, 2800 Plymouth Road Ann Arbor, MI 48109, USA. SF and SV contributed equally to this work.

**Keywords:** (6): Crumbs3, Merlin (NF2), Polarity complex, Epithelial barrier function, Apical actomyosin belt, Rho/ROCK signaling

## Abstract

The gastrointestinal epithelium serves as a critical barrier separating intestinal lumen contents from the underlying tissue environment. Structure and function of the apical junctional complex (AJC), comprising tight and adherens junctions, are essential for establishing and maintaining a polarized and functional epithelial barrier. In this study, we investigated mechanisms by which an apical polarity protein Crumbs homolog 3 (CRB3) regulates AJC assembly and barrier function in primary murine intestinal epithelial cells. Using primary colonic epithelial cells (colonoids) derived from inducible and conditional Crb3 knockdown (Crb3^ERΔIEC^) and control mice (Crb3^fl/fl^), we demonstrate that Crb3 loss leads to compromised epithelial barrier function that was associated with hypercontractile perijunctional actomyosin and defective assembly of the AJC. We identified CRB3 associates with the Band 4.1 family of cytoskeletal linker proteins, Merlin (NF2) via FERM (band4.1/ezrin/radixin/moesin) binding domain (FBD) of CRB3. Interestingly, NF2 knockdown in cultured intestinal epithelial cells phenocopied the effect of CRB3 deletion, supporting a coordinated role in AJC formation and barrier assembly. Moreover, increased active Rho was detected in assembling junctions of Crb3-null cells and inhibition of ROCKII and myosin II alleviated the hypercontractile phenotype, highlighting involvement of Rho/ROCK signaling. Additionally, increased vinculin localization at the AJC seen in Crb3-null epithelial cells indicates elevated tension at junctions. Our findings underscore the important role of Crb3 and NF2 in regulating contractility of the perijunctional actomyosin ring, mechanical tension at the AJC and barrier function via Rho/ROCK signaling during junctional assembly in intestinal epithelial cells.

## Introduction

Epithelial cells reside at the interface of distinct environments, playing crucial roles in maintaining barrier function. In the gut, a monolayer of epithelial cells forms a selective barrier that protects underlying tissues from antigens and pathogens while selectively regulating the movement of nutrients, ion, and fluids. This critical epithelial barrier function is controlled by intercellular junctions, including apical tight junctions (TJs) and adheres junctions (AJs), which are collectively referred to as the Apical Junctional Complex (AJC). The assembly and maintenance of proteins within the AJC are regulated by signaling proteins, mechanical forces, and the interaction of junctional proteins with the underlying perijunctional actin cytoskeleton^1,2^. Thus, failure to assemble or maintain the AJC compromises the epithelial barrier function, which has been implicated in the pathogenesis of chronic inflammatory mucosal diseases such as Inflammatory Bowel Diseases (IBD). Given a critical role of the AJC in controlling epithelial barrier function, it is important to understand mechanisms that govern its formation and maintenance in organ systems such as the intestine.

The Crumbs (CRB) family of proteins play an important role in maintaining apical-basal polarity and regulating intercellular junctions^3–5^, organization of perijunctional actin cytoskeleton^6^, cell proliferation^7,8^ , and morphogenesis in epithelial tissues^9,10^. Three Crumbs paralogs (Crb1, Crb2 and Crb3) are expressed in different tissues. Among these, Crumbs 3 (CRB3) is the most widely expressed paralog and is abundant in the mammalian intestinal epithelial cells. Recent studies have shown that mammalian CRB3 is localized in the apical membrane and the apical vertebrate marginal zone (VMZ) of TJs^11^. Additionally, several *in vitro* studies have revealed that loss of Crb3 results in defects in cell polarity and impaired epithelial junctions, contributing to tumor progression and metastasis in epithelial tissues.

Crb3 is a transmembrane protein that has an extracellular domain, a single transmembrane domain, and a short intracellular domain. The intracellular domain of Crb3 contains a juxtamembrane FERM (band 4.1/ezrin/radixin/moesin) binding domain (FBD) and a carboxy-terminal PDZ (PSD-95/Dlg/ZO1) binding motif. The PDZ binding motif of Crb3 interacts with other polarity complex proteins, including Protein Associated with Lin Seven (Pals1) and Pals1 Associated Tight Junction protein (Patj), PAR, and aPKC via Pals/Patj/PAR/aPKC complex^5,12,13^. Crb3 interaction with Pals/Patj/PAR/aPKC complex has a well-established role in determining apical-basal polarity in epithelial cells^14,15^. While previous studies using transformed epithelial cell lines have suggested a potential role for CRB3 in regulating barrier function, its specific role and underlying mechanisms in AJC regulation in primary mammalian intestinal epithelial cells remain undefined. Furthermore, the contribution of the Crb3 FERM binding domain to regulation of intercellular junctions and epithelial barrier function has yet to be explored.

In this study, we investigated how CRB3 regulates AJC assembly and barrier function in primary murine intestinal epithelial cells. We generated mice with conditional tissue-specific knockdown of CRB3 in the intestinal epithelium. These mice were used to establish primary colonic epithelial cultures, referred to as colonoids. The knockdown of CRB3 led to compromised epithelial barrier function during junctional assembly and was associated with disorganization of the perijunctional actomyosin belt, and mislocalization of AJC proteins ZO-1 and E-cadherin. We identified that CRB3 interacts in a protein complex with the Band 4.1 family of cytoskeletal linker proteins, merlin (NF2) via FERM (band4.1/ezrin/radixin/moesin) binding domain (FBD) of CRB3. Interestingly, knockdown of NF2 in cultured intestinal epithelial cells phenocopied the effect of CRB3 loss on perijunctional actomyosin architecture and barrier integrity. Using primary Crb3 knockdown cells, we demonstrated that reintroducing full-length CRB3a can restore localization of AJC proteins and organization of perijunctional actomyosin. However, this rescue was not achieved with CRB3a mutants lacking either PDZ or FERM binding domains, highlighting a crucial role of these domains play in CRB3 regulation of AJC assembly. Mechanistically, these changes were associated with increased spatial activation of RhoA at the AJC and the formation of a hypercontractile perijunctional actomyosin ring, mediated by Rho/ROCK signaling. In summary, our findings underscore the important roles of CRB3 and NF2 in orchestrating perijunctional actomyosin organization, assembling the AJC, and barrier function in colonoids.

## RESULTS

### Expression of Crb3 in the apical junctional complex of intestinal epithelial cells and generation of *Villin-Cre-ER^T^*^2^ inducible Crb3 knockout mice (Crb3^ΔIEC^)

Previous studies have reported that Crb3 regulates intestinal epithelial morphogenesis. Germline deletion of Crb3 leads to neonatal lethality in mice. To further understand the function of Crb3 in intestinal epithelial cells, we immunolocalized Crb3 in colonic tissue section of a healthy human (Figure 1a) and mouse colon (Figure 1b). As shown in figure 1a, Crb3 localizes in the apical membrane and at the apical junctional complex (AJC) in contrast to E-cadherin that spans the lateral membrane in human colonic epithelial cells. Furthermore, immunolabeling of mouse colonic tissue confirmed analogous Crb3 localization in murine epithelial cells (Figure 1b).

**Figure 1.**
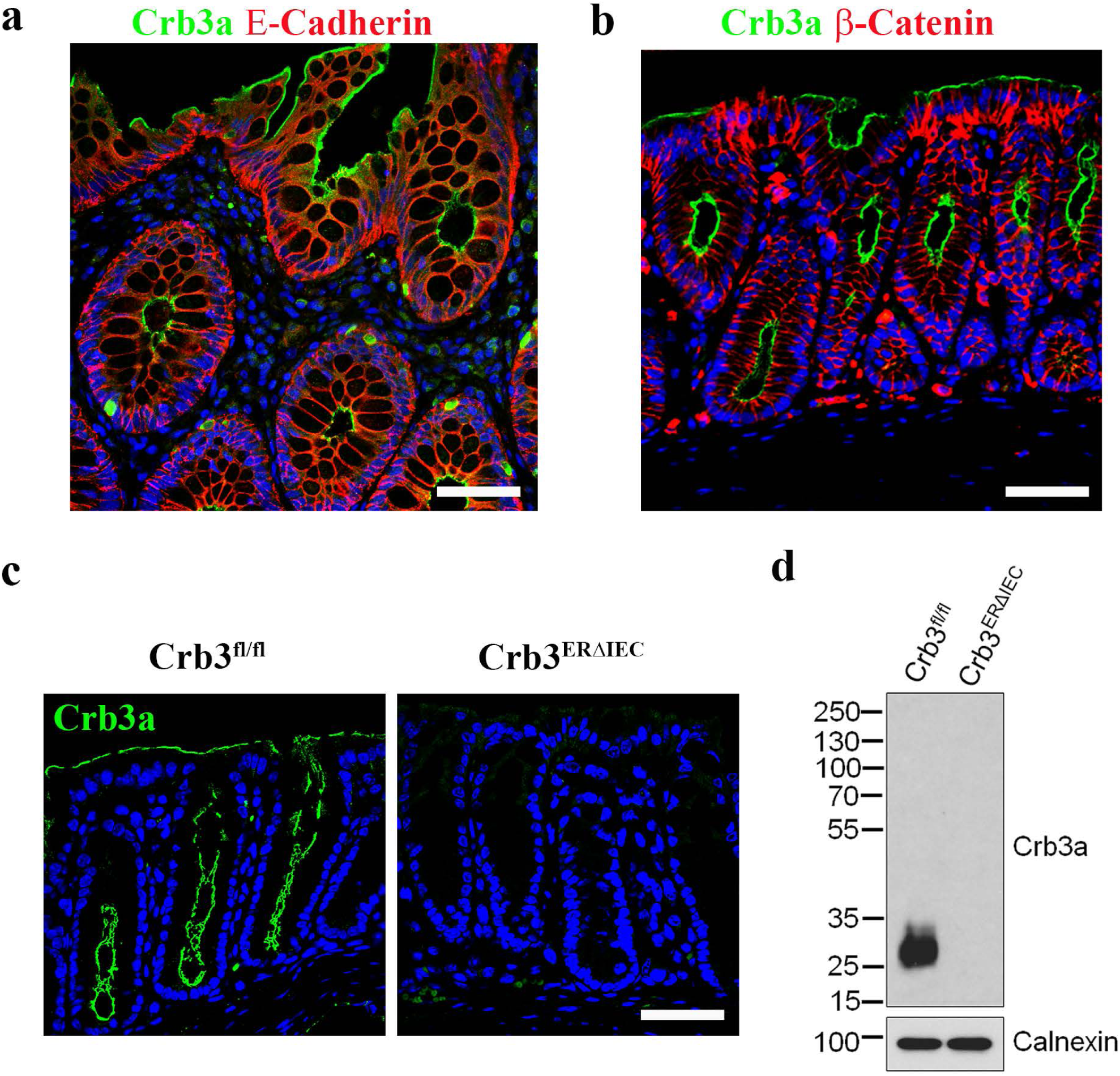
Inducible intestinal epithelial knockout of Crb3 (Crb3 ^ΔIEC^). **(a)** Colonic tissue sections of human healthy donor were immunostained for Crb3a (green), E-Cadherin (red), and DAPI (nuclei/blue). Scale bar = 50 µm; **(b)** Confocal image of murine colonic tissue sections immunostained for Crb3a (green), β-catenin (red), and DAPI (nuclei/blue). Scale bar = 50 µm; **(c)** Immunofluorescence labeling of colonic tissue sections of tamoxifen-treated *Crb3^fl/fl^* and *Crb3^ERΔIEC^* mice showing Crb3a (green) and DAPI (nuclei/blue). Scale bar = 50 µm; **(d)** Immunoblotting of Crb3a and Calnexin (loading control) in colonoids harvested from tamoxifen-treated *Crb3^fl/fl^* and *Crb3^ERΔIEC^* mice. Molecular weight markers are displayed on the left.

Considering the pivotal role of the apical junctional complex in regulating epithelial barrier function, we investigated the impact of Crb3 knockout in epithelial cells on barrier properties. Mice with tamoxifen-inducible deletion of CRB3 specifically in intestinal epithelial cells (Crb3^ERΔIEC^) were generated by crossing Crb3^fl/fl^ mice with intestinal cre reporter Villin-Cre-ER^T^^2^ mice^9^. The efficiency of Crb3 knockout in colonoids was confirmed by immunofluorescence labeling of the murine colonic mucosa (Figure 1c). Additionally, Crb3 knockout was confirmed by immunoblotting, using cells harvested from Crb3^ERΔIEC^ mice and comparing them to Crb3^fl/fl^ matched littermates (Figure 1d). Immunoblotting using an antibody that recognizes all 3 isoforms of Crumbs (Crb1, Crb2 and Crb3) further shows that Crb3 is the only Crb expressed in murine intestinal epithelial cells based on the molecular weight^9^ (Figure 1d)). Off note, the colonic mucosa of the Crb3^ERΔIEC^ mice had normal architecture and was indistinguishable from that of littermate Crb3^fl/fl^ mice, as shown by immunofluorescence labeling/confocal microscopy and histology of the colon (Supplemental Figure 1a-b). Collectively, our results demonstrate efficient Crb3 knockdown in Crb3^ERΔIEC^ intestinal epithelial cells.

### Intestinal epithelial Crb3 regulates localization of AJC proteins, organization of perijunctional actin cytoskeleton and barrier function

The interaction of Crb3 with Pals1/Patj and Par3/Par6/aPKC protein complexes in epithelial cells is important for regulating the apical junctional complex (AJC), which includes the tight junction (TJ) and adherens junction (AJ). Immunostaining and confocal imaging of sub-confluent colonoids with assembling intercellular junctions harvested from Crb3^fl/fl^ and Crb3^ERΔIEC^ mice revealed discontinuous junctional localization of Patj and Par3 in the absence of Crb3 (Figure 2a). This finding corroborates previous reports, demonstrating that Crb3 stabilizes its binding partners Patj and Pals1 at cell-cell junctions. Additionally, Par3 and the TJ scaffold protein, ZO-1, exhibited discontinuous labeling and reduced intensity at TJs of colonoids in the absence of Crb3 (Figure 2a and 2c). E-cadherin was visualized as linear staining in the junction plasma membrane of epithelial cells derived from Crb3^fl/fl^ mice. However, in epithelial cells lacking Crb3 (Crb3^ERΔIEC^) E-cadherin was visualized as diffuse and disrupted staining in the lateral plasma membrane (Figure 2a). Additionally, immunoblotting of colonoids derived from Crb3^fl/fl^ and Crb3^ERΔIEC^ mice revealed decreased Crb3 binding partner proteins (Patj, Pals1, and Par3) and decreased tight junction proteins (TJ, ZO-1, ZO-3) in cells lacking Crb3 (Crb3^ERΔIEC^) (Figure 2b). However, the expression level of adherens junction (AJ) protein, E-Cadherin was unchanged in colonoids in the absence of Crb3 (Figure 2b). These results suggest that Crb3 regulates the localization and organization of AJC proteins during junction assembly in intestinal epithelial cells.

**Figure 2.**
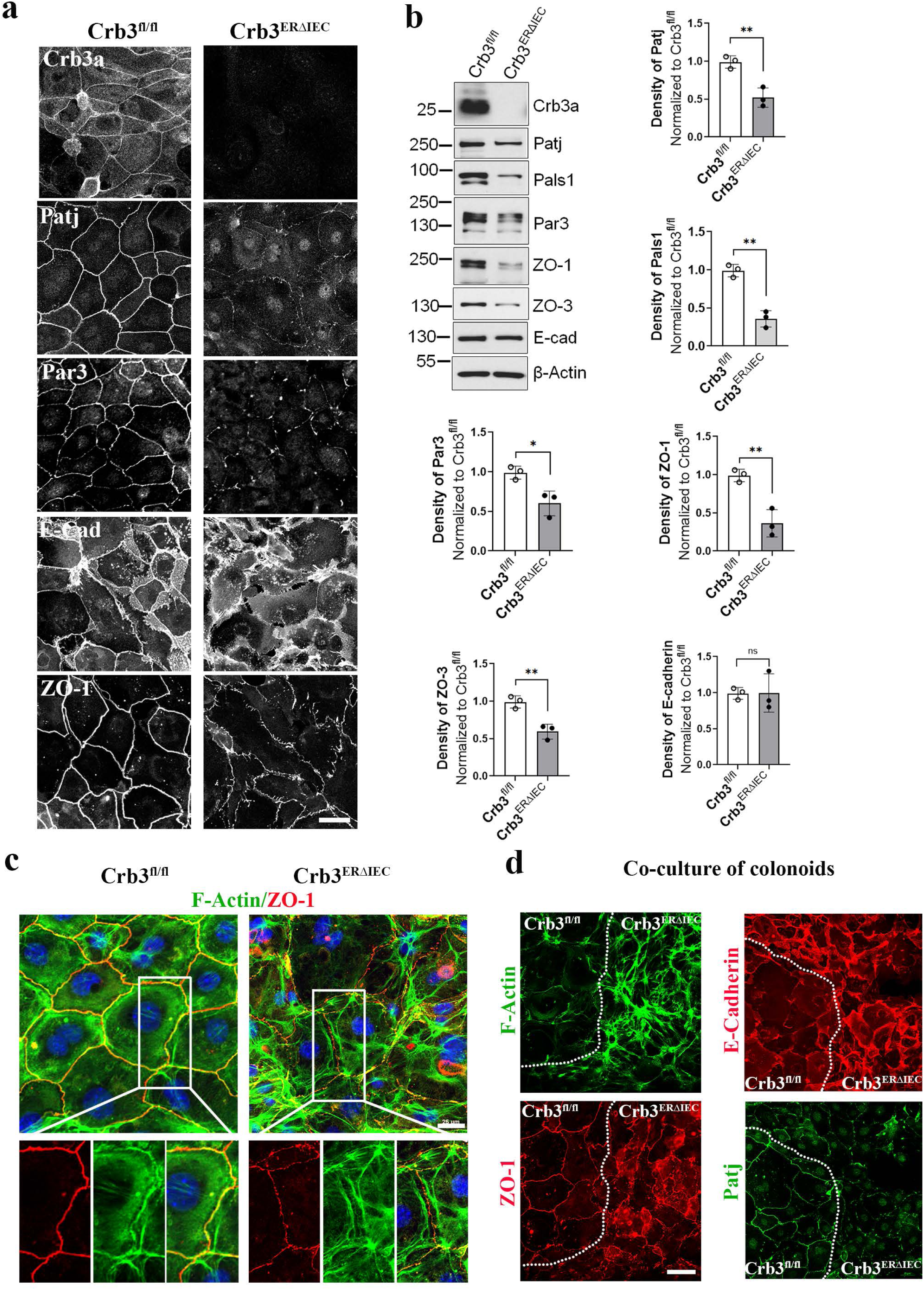
Crb3 regulates junctional localization of AJC proteins and organization of perijunctional actin cytoskeleton in intestinal epithelial cells. **(a)** Representative confocal images of colonoids derived from Crb3^fl/fl^ and Crb3^ERΔIEC^ mice during junctional assembly (subconfluent) immunostained for polarity proteins (Crb3, Patj, Par3), AJ protein (E-cadherin), and the TJ protein (ZO-1); **(b)** Immunoblotting of colonoids derived from Crb3^fl/fl^ and Crb3^ERΔIEC^ mice during junctional assembly (subconfluent) for polarity proteins (Crb3, Patj, Pals1, Par3), AJ protein (E-cadherin), TJ protein (ZO-1, ZO-3), and loading control (β-actin).Graph showing the quantification of immunoblots in 2b. Data are mean ±SD, n=3 experiments. **p≤0.01; ***p≤0.001 by two tailed students t test; **(c)** Representative confocal images of F-Actin (Phalloidin, green), TJ protein (ZO-1, red), and nuclei (DAPI, blue) in colonoids derived from Crb3^fl/fl^ and Crb3^ERΔIEC^ mice. White rectangles represent the zoom-in areas shown below the composite images. Scale bar = 25µm; **(d)** Coculture of colonoids derived from Crb3^fl/fl^ and Crb3^ERΔIEC^ mice stained for F-Actin (Phalloidin, green), ZO-1 (red), Patj (green), and E-Cadherin (red). White dotted line marks the boundary between cells expressing Crb3 and cells lacking Crb3. Scale bar = 25µm.

In epithelia, organization and contractility of the apical perijunctional actin cytoskeleton are crucial for the formation and maintenance of the AJC and therefore epithelial barrier function^16–18^. Since Crb3 expression influenced the junctional localization of AJC proteins, we determined if Crb3 expression modulates perijunctional actin cytoskeleton. Co-labeling of ZO-1 and F-actin in colonoids from Crb3^fl/fl^ and Crb3^ERΔIEC^ mice revealed significant alterations in the perijunctional F-actin of Crb3^ERΔIEC^ epithelial cells, which displayed a “railroad” appearance along the AJC compared to the control Crb3^fl/fl^ epithelial cells. In Crb3 knockout cells, this F-actin organization suggested a hypercontractile state (Figure 2c). To minimize interpretation bias, we co-cultured colonic epithelial cells derived from Crb3^fl/fl^ and Crb3^ERΔIEC^ mice at 1:1 ratio and labeled for F-actin, Patj, ZO-1, and E-Cadherin. Based on F-actin morphology and disrupted ZO-1 staining, we delineated the boundary between Crb-expressing cells (left) and Crb3 knockout epithelial cells (right) with a white dotted line in Figure 2d. In Crb3 knockout epithelial cells, ZO-1 and E-cadherin were mislocalized in the AJC which was associated with disorganization of perijunctional F-actin. Consistent with Figure 2a, the junctional location of Crb3 binding partner protein, Patj, was significantly reduced in Crb3 knockout colonic epithelial cells (Figure 2d). Together, these results support the role of Crb3 in regulating the junctional localization of AJC proteins and the organization of perijunctional F-actin cytoskeleton during junctional assembly in intestinal epithelial cells.

Defects in formation and maintenance of intestinal epithelial barrier are strongly associated with altered expression and localization of AJC proteins, including ZO, E-Cadherin, Pals1 and Patj. Since loss of Crb3 leads to disorganization of the AJC, we evaluated epithelial barrier function by measuring the transepithelial electrical resistance (TER) and paracellular flux of solute. Many previous studies have relied on immortalized or cancer cell lines to either overexpress or transiently knockdown Crb3. We, therefore, analyzed the role of Crb3 on epithelial barrier assembly using colonoids that were harvested from Crb3^fl/fl^ and Crb3^ERΔIEC^ mice and cultured as monolayers on permeable supports. As shown in figure 3a, TER of epithelial cells from Crb3^fl/fl^ increased substantially over 5 days which is associated with junctional assembly and maturation. However, TER of epithelial cells from Crb3^ERΔIEC^ mice was significantly lower compared to Crb3^fl/fl^. Additionally, increased paracellular flux of 4kDa dextran was observed in concert with the decreased TER in Crb3^ERΔIEC^ epithelial cells compared to the control cells (Figure 3b). These results collectively suggest that Crb3 plays an important role in promoting the formation of the intestinal epithelial barrier by regulating the junctional localization of AJC proteins and organization of perijunctional F-actin cytoskeleton.

**Figure 3.**
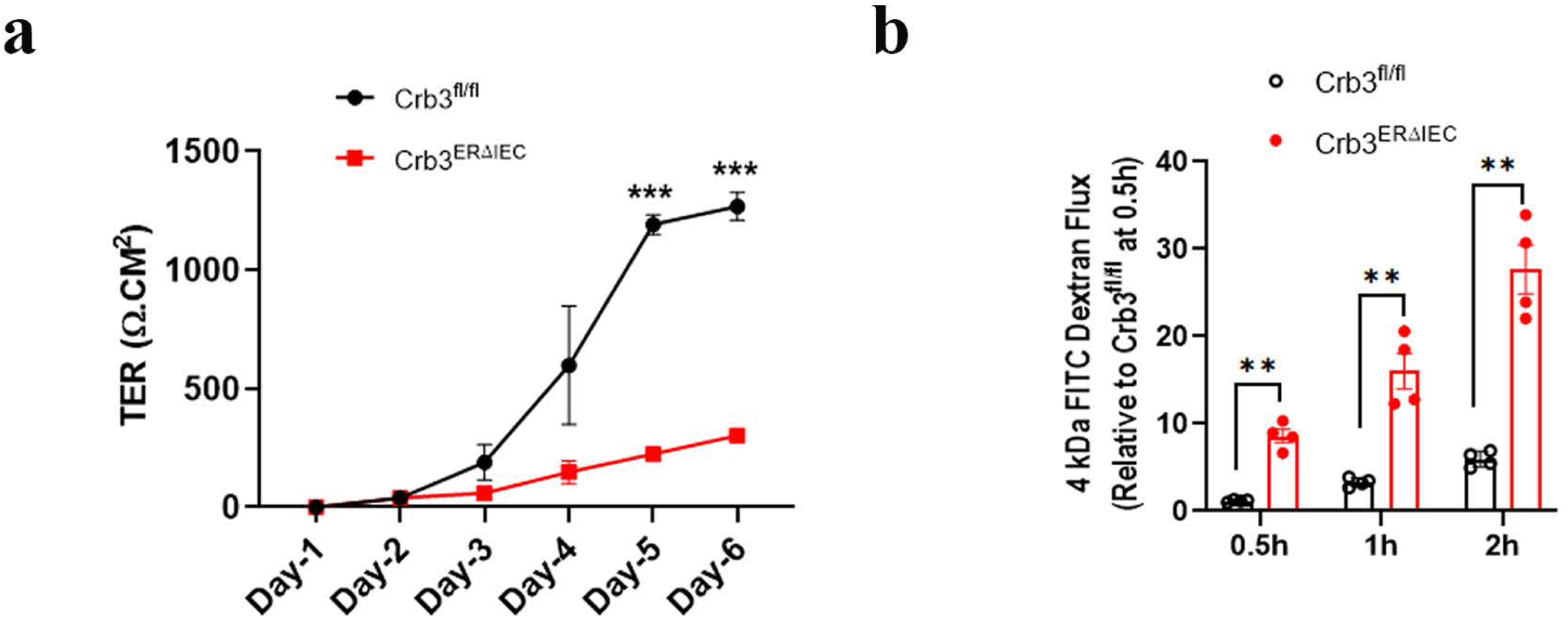
Crb3 regulates the formation of the intestinal epithelial barrier. **(a)** Graph showing establishment of epithelial barrier function that was measured by TER in confluent colonoids from Crb3^fl/fl^ (shown in black) and Crb3^ERΔIEC^ (shown in red) mice grown cultured on transwells for 6 days (Day 1 represents day of isolation). Data are Mean±SD, n=3 experiments each with six technical replicates. *** p≤0.001 by two tailed students t test; **(b)** Paracellular flux rate of 4kDa FITC-dextran across monolayers of primary murine colonoids derived from Crb3^fl/fl^ and Crb3^ERΔIEC^ mice at times indicated in the x-axis. Data are mean ± SD. n=4 experiments each with six technical replicates ** p≤0.01 by two tailed students t test.

### PDZ binding motif and FERM binding domain (FBD) of Crb3a regulates assembly of the AJC and perijunctional F-actin ring in intestinal epithelial cells

Crb3a C-terminal cytoplasmic tail contains a FERM binding domain (FBD) and PDZ binding motif (PBM). In vertebrates, Crb3a is known to associate with Ezrin, a member of Ezrin-Radixin-Moesin (ERM) protein family through FBD^9^. ERM proteins have been reported to play a crucial role in organizing the perijunctional actomyosin ring by anchoring it to the cell-cell junctions in epithelial cells^19,20^. To elucidate the mechanism by which Crb3a controls AJC assembly and barrier function, we assessed the roles of Crb3a FBD and PBM in regulating these processes in primary Crb3^ERΔIEC^ colonic epithelial cells. We cloned wildtype and mutant human Crb3a tagged with an N-terminal Myc epitope into a pLentiviral vector (pLLV). Myc tagged Crb3a wildtype (WT), FBD mutant, or ΔPDZ mutants were expressed in primary mouse colonoids isolated from Crb3^ERΔIEC^ mice. The schematic in Figure 4a illustrates the constructs: wild-type human Crb3a (pLLV myc-Crb3 wt), Crb3a mutant with three amino acid point mutations in FBD (pLLV myc-FBD mut, highlighted in red), and Crb3a mutant lacking ERLI sequence (pLLV myc-Crb3 ΔPDZ).

**Figure 4.**
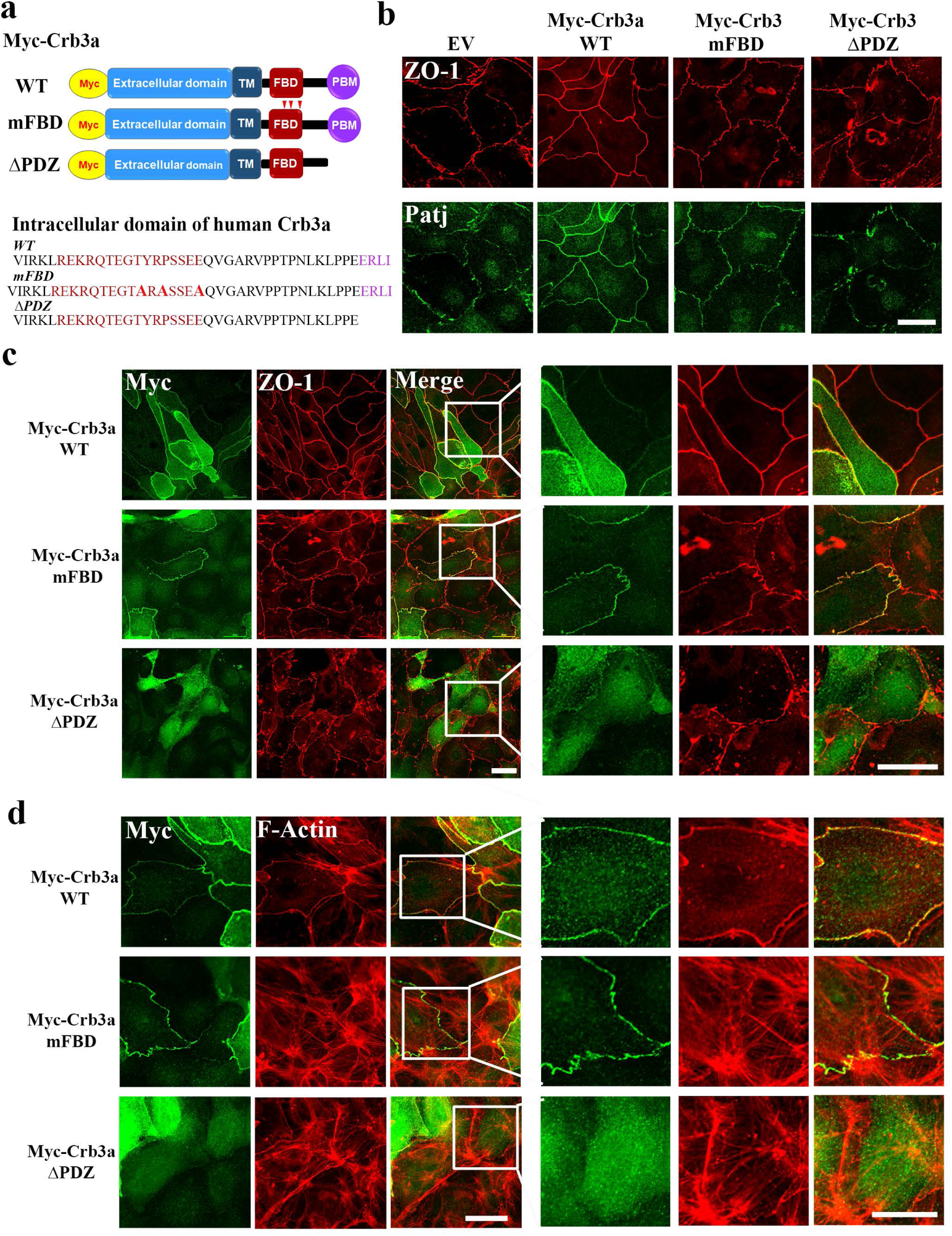
PDZ binding motif and FERM binding domain (FBD) of Crb3a control assembly of the AJC and perijunctional F-actin ring in intestinal epithelial cells. **(a)** Schematic of wild-type human Crb3a (Crb3 wt), Crb3a mutant with 3 amino acid point mutations in FBD (FBD mut, red arrows) and Crb3a mutant lacking PDZ binding motif (PBM, ΔPDZ). Amino acids in FBD shown in brown with mutations shown in red and PBM in purple; **(b)** Immunofluorescence labeling and confocal images of apical junctional proteins, ZO-1 (red, upper panel) and Patj (green, lower panels), during junction assembly in sub-confluent colonoids derived from *Crb3^ERΔIEC^* mice infected with lentiviral Crb3 wt (pLLV myc-Crb3a wt), ΔPDZ motif (pLLV myc-Crb3a ΔPDZ), and FBD mut (pLLV myc-Crb3a FBD mut) constructs; **(c)** Co-staining of apical junction protein, ZO-1 (red) and myc-tag (anti-myc, green) in sub-confluent colonoids derived from *Crb3^ERΔIEC^*mice expressing plasmids described in Fig. 4b. Cells stained positive for Myc (green) represent the Crb3 KO murine colonoids expressing the lentiviral Crb3a constructs described in Fig. 4a. White box of the zoomed area shown on the right panel. Scale bar = 25 µm; **(d)** Co-staining of perijunctional F-actin ring (Phalloidin, red) and myc-tag (green) in colonoids derived from *Crb3^ERΔIEC^* mice described in Fig. 4b. White box highlights the zoomed area shown in the right panel. Scale bar = 25 µm.

Colonoids harvested from Crb3^ERΔIEC^ mice were transduced with lentivirus to express Crb3a WT (pLLV myc-Crb3a wt), Crb3a ΔPDZ mutant (pLLV myc-Crb3a ΔPBM), or Crb3a FBD mutant (pLLV myc-Crb3a mFBD). Immunofluorescence labeling for ZO-1 (red, upper panel) and Patj (green, lower panels), revealed that both Crb3a FBD mut and Crb3a ΔPDZ mutants were unable to restore the linear junctional plasma membrane localization of ZO-1 and Patj, in contrast to the expression of Crb3 WT (Figure 4b). Next, we performed co-immunostaining of myc, ZO-1 (Figure 4c) and F-actin (Figure 4d) in colonoids from Crb3^ERΔIEC^ mice expressing Crb3a WT or Crb3a mutants. Myc immunostaining demonstrated junctional localization of Crb3a in Crb3-null colonoids expressing either Crb3a WT or Crb3a FBD mutant (Figure 4c). However, the Crb3a ΔPDZ mutant failed to localize in the junctional plasma membrane in Crb3-null colonoids (Figure 4c) indicating that Crb3a PDZ is required for Crb3a targeting to the epithelial AJC. Furthermore, expression of myc-Crb3a WT in Crb3-null colonoids rescued the junctional localization of ZO-1 (Figure 4c) and perijunctional F-actin organization (F-actin, Figure 4d), while Crb3a mutants (myc- Crb3a mFBD and myc-Crb3a ΔPDZ) failed to rescue these parameters (Figure 4c, d). Collectively, our results demonstrate that the Crb3a PBM is crucial for the junctional localization of Crb3a. Moreover, both Crb3a FDB and PBM play pivotal roles in regulating apical junctional assembly and the organization of the perijunctional F-actin ring in intestinal epithelial cells.

### NF2 (Merlin) interacts with FERM binding domain (FBD) of Crb3a in colonoids

Our results demonstrate that cytoplasmic FBD of Crb3a regulates TJ assembly and organization of the perijunctional F-actin ring. In Drosophila, Crb interacts with Expanded (Ex), and two other FERM-containing proteins, Yurt and DMoesin, via the Crb cytoplasmic FBD to regulate the Hippo signaling pathway. In humans, Crb3, the most widely expressed ortholog binds to Ezrin and regulates morphogenesis of intestinal villi, and microvilli. In intestinal epithelial cells, ezrin exclusively localizes apical to the TJ (ZO-1) in the vertical marginal zone (VMZ), which is the apical-most region of the lateral plasma membrane. Given the apical localization of ezrin which is not within the AJC, it is likely that other members of the ERM protein family interact with colonic epithelial Crb3 to regulate AJC assembly and the organization of perijunctional F-actin.

NF2 (Merlin), a member of the ERM family, is known to localize to the AJC and interact with a TJ protein Angiomotin^21^, and AJ protein α-Catenin in epithelial cells^20^. To investigate whether Crb3a is a part of a protein complex with NF2, we evaluated the potential association between Crb3a and NF2. Co-immunoprecipitation of Crb3a with NF2 was performed using a model intestinal epithelial cell line (SKCO-15) expressing myc tagged full length Crb3a (myc- Crb3a WT). As expected, anti-myc antibodies co-immunoprecipitated with endogenous NF2 (Figure 5a). Conversely, endogenous NF2 co-immunoprecipitated with endogenous Crb3a in SKCO-15 cells (Figure 5b). Expression of phosphorylated NF2S518, which is inactive for its ability to have tumor suppression function^22^ , failed to pull down Crb3a in SKCO-15 cells (Figure 5b). Furthermore, expression of phosphorylated NF2 at serine 518 (NF2S518), which is inactive, failed to pull down Crb3a in SKCO-15 cells (Figure 5b). Additionally, immunostaining of sub-confluent colonic epithelial cells from Crb3^fl/fl^ and Crb3^ERΔIEC^ mice with assembling junctions demonstrated that NF2 localization was not confined to the AJC but extended across the entire lateral membrane in Crb3^ERΔIEC^ epithelial cells, compared to control Crb3^fl/fl^ cells where NF2 co-localized with ZO-1 (Figure 5c). However, expression levels of NF2 were not altered in colonic epithelial cells from either Crb3^fl/fl^ and Crb3^ERΔIEC^ mice (Supplemental Figure 2).

**Figure 5.**
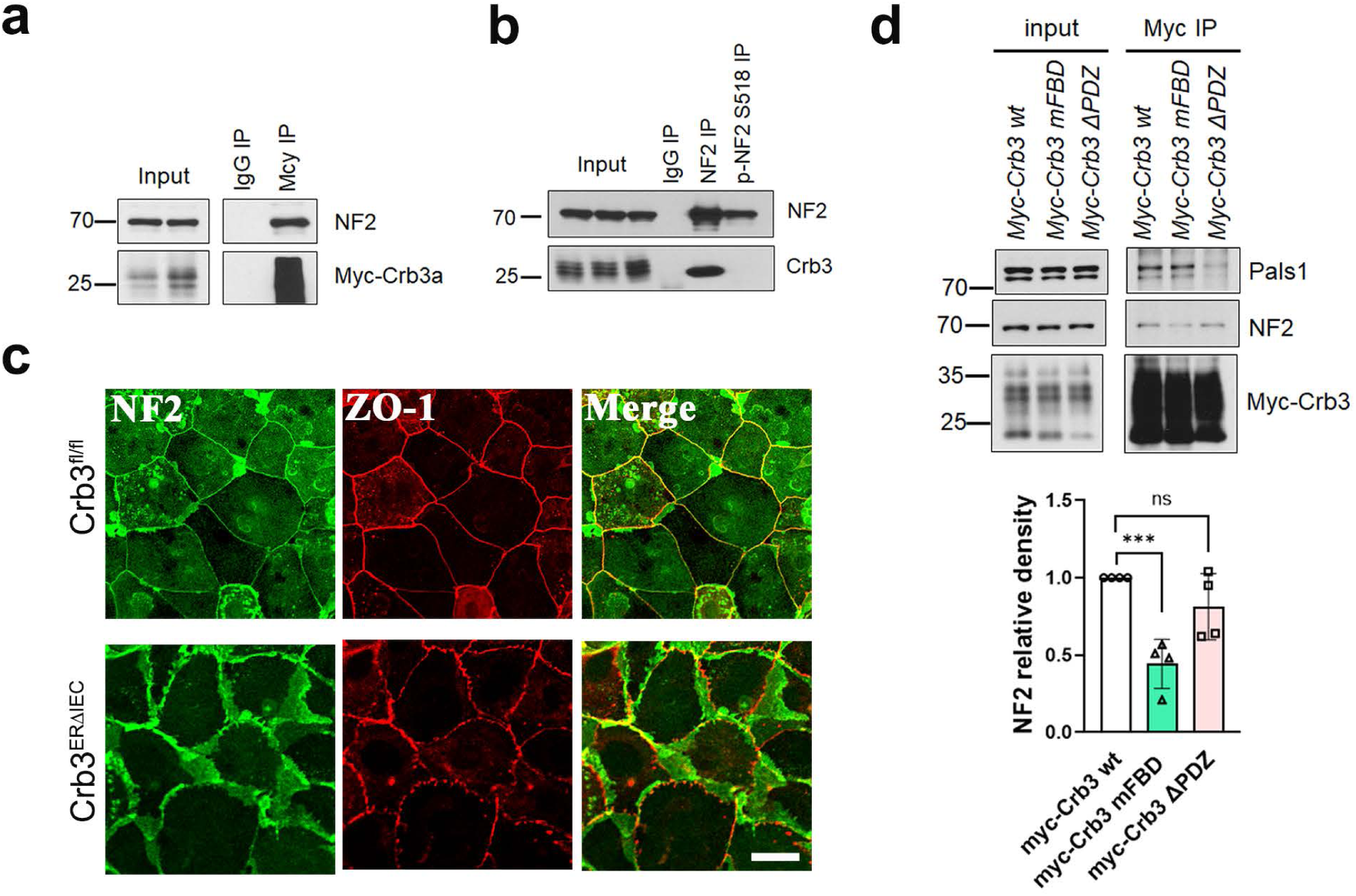
NF2 interacts with Crb3a in intestinal epithelial cells and regulates epithelial junction assembly. **(a)** Immunoprecipitation of Crb3a in SK-CO15 IECs expressing Myc-Crb3a wt followed by western blotting for NF2 and Myc- tag. Myc-Crb3 immunoprecipitates NF2. IgG immunoprecipitation serves as a negative control; **(b)** Immunoprecipitation of NF2 and p-NF2 (S518) in SK-CO15 IECs followed by western blotting for Crb3a; **(c)** Representative confocal images of immunostained proteins showing the junctional localization of NF2 (green), ZO-1 (red), and merge in subconfluent murine colonoids derived from Crb3^fl/fl^ and Crb3^ERΔIEC^ mice. Scale bar = 25 µm; **(d)** (Top) Immunoprecipitation of anti-Myc antibody in SK-CO15 IECs expressing Myc- Crb3a wt, Crb3a FBD mut, or Crb3a ΔPDZ followed by western blotting for NF2, Pals1 and Myc-tag. Decreased binding of NF2 in Crb3a FBD mut, while Crb3a ΔPDZ did not associate with Pals1. (Bottom) Graph showing densitometry of NF2 immunoblot relative to loading control and normalized to SK-CO15 IECs expressing Myc-Crb3a wt. Data are mean ± SD. n=4. Each dot represents an independent experiment *** p≤0.001 by two tailed students t-test.

Next, we assessed the interaction of NF2 with the FBM and PDZ binding motif of Crb3a by performing co-immunoprecipitation experiments in SKCO-15 cells expressing myc-tagged Crb3a WT and myc-tagged Crb3a mutants. Our results indicate that while anti-myc antibodies co-immunoprecipitated endogenous NF2 with myc-Crb3a WT and myc-Crb3a ΔPDZ, the interaction was significantly reduced with myc-Crb3a mFBD (Figure 5d). Additionally, the interaction between Crb3a and Pals1 was reduced only in cells expressing myc-Crb3a ΔPDZ mutant, consistent with previous studies^3^ (Figure 5d). These results collectively demonstrate that NF2 interacts with FBD of Crb3a, and that the junctional localization of NF2 is dependent on Crb3a.

### NF2 knockdown phenocopies the effects of Crb3 deletion, disrupting AJC assembly and compromising epithelial barrier function in intestinal epithelial cells

NF2 plays an important role in anchoring the perijunctional actomyosin ring to the plasma membrane^20^. Based on our results showing that Crb3 interacts with NF2 in intestinal epithelial cells, we investigated whether NF2 has a similar role to Crb3 in AJC assembly, the organization of the perijunctional F-actin ring, and the epithelial barrier function.

NF2 was knocked down (KD) in model intestinal epithelial cells, SKCO-15 using two independent siRNAs targeting NF2. As shown in Figure 6a and Supplemental Figure 3, both NF2 siRNAs downregulated NF2 compared to scrambled control siRNA. Barrier function was analyzed by measuring TER of epithelial cells cultured on permeable supports over a period of 4 days. Significant decrease in TER development was observed in NF2 KD epithelial cells compared to control epithelial cells (Figure 6 a and b). Co-immunostaining of F-actin and ZO-1 in control and NF2 KD cells revealed that NF2 knockdown led to disrupted organization of the perijunctional F-actin ring and discontinuous ZO-1 staining at the TJ, similar to the effects observed in epithelial cells lacking Crb3 (Figure 6c).

**Figure 6.**
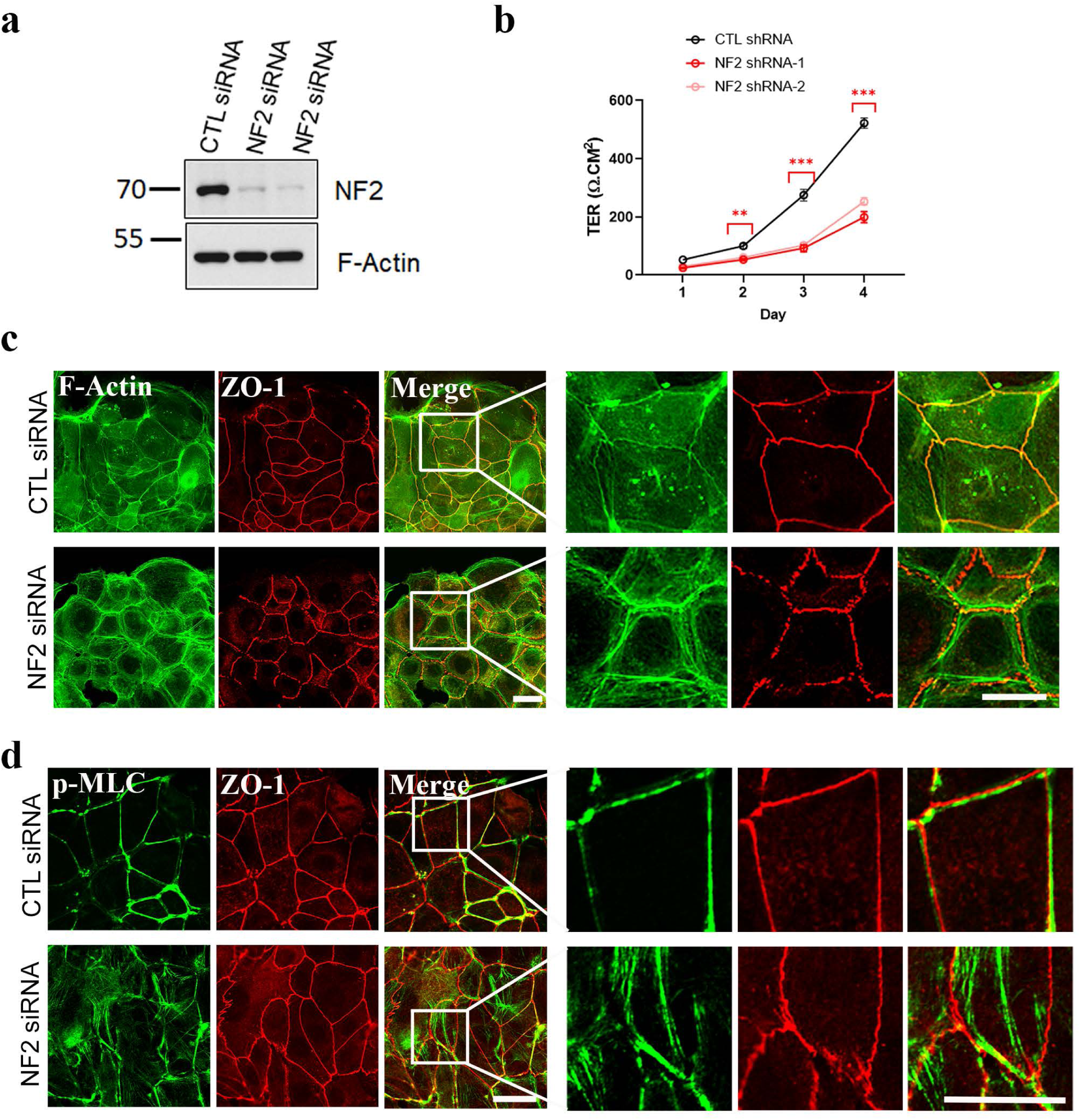
Loss of NF2 mimics the defects seen in Crb3-deficient IECs, including impaired apical junction assembly, disorganization of the perijunctional actomyosin belt, and compromised epithelial barrier function. **(a)** Representative immunoblot showing the efficiency of NF2 knockdown in SK CO-15 IECs transfected with control scramble or NF2 siRNA with F-Actin as a loading control; **(b)** Graph showing TER of SKCO15 cells with siRNA mediated knockdown of NF2 or scramble control in cells cultured as monolayers for 4 days. Data are mean ±SD, n=3 experiments each with six technical replicates. **p≤0.01; ***p≤0.001 by two tailed students t-test; **(c)** Representative perijunctional F-actin organization (Phalloidin, green) and tight junction protein ZO-1 (red) localization in sub confluent SK CO-15 cells with siRNA knockdown of NF2 or control scramble siRNA for 48 hours. White box highlights an area that is enlarged in the right panel. Scale bar = 25 µm; **(d)** Immunostaining of junctional myosin (p-MLC T18/S19, green) and the tight junction protein ZO-1 (red) in sub confluent SK CO-15 with NF2 siRNA mediated knockdown or control scramble siRNA for 48 hours. White box highlights the zoomed area shown in the right panel. Scale bar = 25 µm.

In addition to the organization of the perijunctional F-actin ring, contractility of the actomyosin ring regulates AJC assembly and epithelial barrier function^1,18^. Phosphorylation of the myosin regulatory light chain at threonine 18 and serine 19 (pMLCIII^T^^18/S^^19^) controls perijunctional actomyosin contraction. Co-immunostaining for pMLCIII^T^^18/S^^19^ and ZO-1 in NF2 knockdown cells revealed disorganization and displacement of pMLCII from the AJC compared to control cells, indicative of a hypercontractile perijunctional actomyosin ring (Figure 6d). Together, our results demonstrate that NF2 regulates AJC formation and the establishment of epithelial barrier function, mirroring the role of Crb3 in intestinal epithelial cells.

### Loss of Crb3 results in a hypercontractile perijunctional actomyosin ring, driven by enhanced RhoA-ROCK signaling in intestinal epithelial cells

The small GTPase RhoA, in its active form (RhoA-GTP), is a key regulator of the perijunctional actomyosin ring in epithelial cells^23,24^. Activated RhoA stimulates its downstream effector ROCKII, which generates contractile forces by phosphorylating MLCII. Co-immunostaining of F-actin and pMLCII in sub-confluent colonoids from Crb3^fl/fl^ and Crb3^ERΔIEC^ mice, demonstrated changes in perijunctional F-actin, which exhibited a “rail-road track” appearance, and mislocalization of junctional pMLCII in Crb3^ERΔIEC^ colonoids (Figure 7a). Additionally, co-localization of ZO-1 and pMLCII at TJs was disrupted in Crb3 knockout cells (Figure 7b). These findings implicate a hypercontractile perijunctional actomyosin ring as responsible for defective AJC formation observed in Crb3 null epithelial cells.

**Figure 7.**
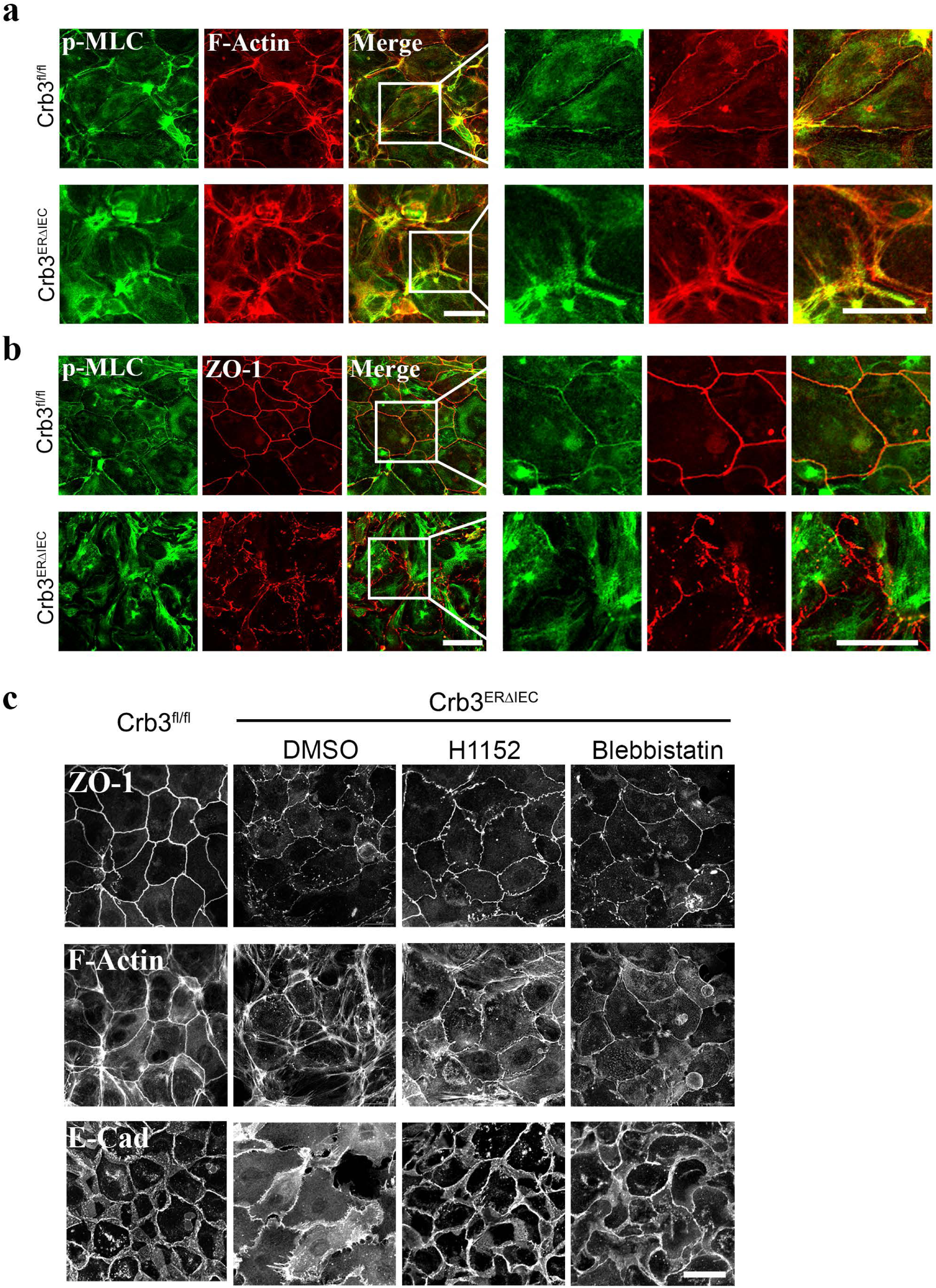
Inhibition of ROCK or p-MLC activity largely restores the defects in apical junction assembly and the organization of the perijunctional actomyosin belt in Crb3 KO IECs. **(a)** Representative confocal images of sub- confluent colonoids derived from Crb3^fl/fl^ and Crb3^ERΔIEC^ mice coimmunostained for p-MLC T18/S19 (green) and F-actin (Phalloidin, red). Scale bar = 25 µm; **(b)** Representative confocal images of sub-confluent colonoids derived from Crb3^fl/fl^ and Crb3^ERΔIEC^ mice coimmunostained for p-MLC T18/S19 (green) and ZO-1 (red). Scale bar = 25 µm; **(c)** Representative confocal images of perijunctional F-actin (Phalloidin), and AJC proteins (ZO-1 and E-Cadherin) in Crb3^fl/fl^ colonoids and Crb3^ERΔIEC^ colonoids treated with ROCK II inhibitor (H1152,10 µM), Myosin II inhibitor (Blebbistatin, 25 µM), or vehicle (DMSO, 0.1%) for 30 minutes. Scale bar = 25 µM.

To investigate this possibility further, sub-confluent colonoids derived from Crb3^ERΔIEC^ mice were treated for 30minutes with ROCK II inhibitors (H1152, and Y27632), a myosin II inhibitor (Blebbistatin), and a vehicle control (DMSO). Immunostaining for ZO-1, E-cadherin, and F-actin demonstrated that inhibiting myosin II contractility with these compounds largely restored general architecture of the perijunctional F-actin ring (Figure 7c and Supplemental Figure 4a) and the junctional localization of AJC proteins including TJ (ZO-1) and AJ (E-cadherin) in Crb3 null IEC (Figure 7c, and Supplemental Figure 4b). Additionally, p-MLC ^T^^18/S1^^9^ staining confirmed that H1152 and Y-27632 suppressed ROCK/MLCP/p-MLC signaling pathway, while Blebbistatin directly inhibited MLC ATP/ADP turnover rather than MLC phosphorylation (Supplemental Figure 4a). Additionally, RhoA activity was analyzed during junctional assembly in colonoids derived from Crb3^fl/fl^ and Crb3^ERΔIEC^ mice. Endogenous levels of active RhoA was examined using a localization-based biosensor for active RhoA (rhoketin G protein binding domain, rGBD). As previously demonstrated, fluorescently tagged rGBD probe (dTomato-2xrGBD) binds specifically to GTP bound state of RhoA (active RhoA)^25,26^ . Live cell imaging of sub-confluent colonoids derived from Crb3^fl/fl^ and Crb3^ERΔIEC^ mice transduced with pLV-dimericTomato-2xrGBD and labeled with live cell membrane probe (MemBright 488), showed increased junctional signal of active Rho in Crb3 null epithelial cells compared to control cells (Figure 8a, b,c). Inhibition of active RhoA with C3 transferase (C3T) in Crb3 null epithelial cells reduced the junctional signal of active RhoA, thereby demonstrating the specificity of dTomato-2xrGBD in detecting active Rho (Supplemental Figure 5c).

**Figure 8.**
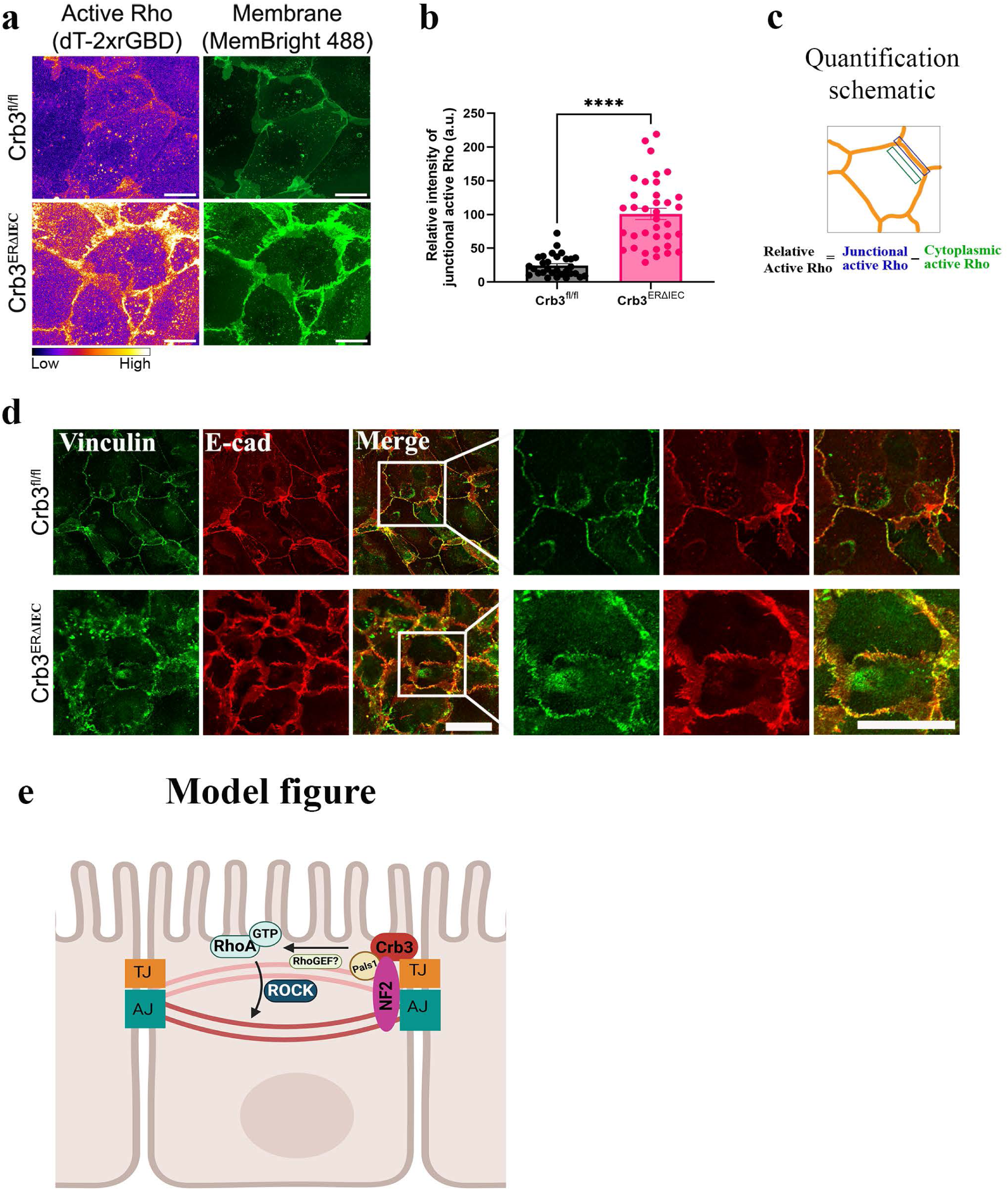
Loss of Crb3 results in increased junctional RhoA activity and increased cell-cell junction tension during junction assembly in IECs. **(a)** Representative airyscan image of sub-confluent colonoids derived from Crb3^fl/fl^ and Crb3^ERΔIEC^ mice expressing pLenti-dTomato-2xrGBD (pseudo colored as FIRE LUT) and stained with MemBright 488 plasma membrane dye (green). Images shown are sum of Z-projections. Scale bar = 20 µm; **(b)** Graph showing the relative intensity of active Rho at cell-cell junctions normalized to the background signal of every junction. Data are mean ± SEM. Each dot represents an independent junction from 3 independent experiments *** p <0.0001 by Mann-Whitney test. n= 30 junctions for Crb3^fl/fl^; n=36 junctions for Crb3^ERΔIEC^; **(c)** Schematic showing the quantification method used to measure the relative intensity of junctional active RhoA; **(d)** Representative confocal images of sub confluent colonoids derived from Crb3^fl/fl^ and Crb3^ERΔIEC^ mice immunostained for Vinculin (green) and E-cadherin (red). White box represents zoomed in region shown on the right. Scale bar = 25 µm; **(e)** Model figure illustrating the role of CRB3 and its binding partner NF2 in regulating perijunctional actomyosin belt and barrier function via Rho/ROCK signaling during junction assembly.

To further evaluate the increased mechanical tension at the AJC in cells lacking Crb3, we examined the localization of vinculin, which is known to be recruited to the AJ by α-Catenin under conditions of increased tension^27–29^. Co-immunostaining of Vinculin and E-Cadherin in sub-confluent colonoids derived from Crb3^fl/fl^ and Crb3^ERΔIEC^ mice, revealed increased vinculin localization at the AJC in Crb3 null epithelial cells compared to control cells (Figure 8d). To demonstrate antibody specificity, two different vinculin antibodies were used for co-immunostaining with ZO-1 and F-actin (Supplemental Figure 5a-b). Together, these results demonstrate that Crb3 expression influences optimal contractility of the perijunctional actomyosin ring and tension at the AJC, mediated by Rho/ROCK signaling during junction assembly in intestinal epithelial cells.

## Discussion

Epithelial morphogenesis is a fundamental process that shapes the structure and function of epithelial tissue during development and tissue homeostasis. Apical cell-cell junctions help establish cell polarity and maintain a functional epithelial barrier, both of which are critical for tissue organization and function. Previous work has shown that CRB3 regulates intestinal epithelial morphogenesis. However, the precise mechanism by which CRB3 regulates intestinal epithelial barrier function in an *in vivo* model or in harvested colonoids is unclear due to premature death of mice with germline deletion of Crb3^9^. In this study, we generated a mouse model with tamoxifen-inducible intestinal epithelial tissue specific deletion of Crb3 (Crb3^ERΔIEC^). The small intestine and colonic mucosa of Crb3^ERΔIEC^ mice displayed normal architecture indistinguishable from that of Crb3^fl/fl^ littermates. This finding suggests that villus fusion and disrupted apical membranes seen in mice with germline mutations of Crb3 result from embryonic organ developmental abnormalities due to a lack of Crb3. Additionally, to analyze the role of CRB3 in junctional assembly, we were able to generate Crb3-null colonoids from Crb3^ERΔIEC^ mice that were compared to wild type counterparts Crb3^fl/fl^, offering insights beyond those obtained from immortalized or cancer cell lines^3,30^. Our results demonstrate that CRB3, regulates the assembly of apical cell-cell junctions and establishment of barrier function in intestinal epithelium.

Establishment of epithelial barrier function requires tightly controlled spatiotemporal assembly of intercellular junctions that include the tight junction, adherens junction and associations with the apical perijunctional actomyosin cytoskeleton. Defects in formation and maintenance of the AJC result in a compromised intestinal epithelial barrier, a hallmark of chronic mucosal inflammatory diseases including Ulcerative Colitis and Crohn’s disease^31^. The assembly and maintenance of apical junctional complexes are influenced by signaling pathways, mechanical forces, and interaction of AJC proteins with the actin cytoskeleton. We identified epithelial barrier compromise in colonoids derived from Crb3^ERΔIEC^ mice that was associated with changes in localization of AJC proteins in the junctional plasma membrane. In addition, there was associated disorganization of the apical perijunctional F-actin ring that exhibited a “rail-road” pattern suggestive of increased actomyosin tension at the apical junctional complex. These changes imply that CRB3 regulates localization of AJC proteins while also playing an important role in organizing the actin cytoskeleton, an essential element in the control of epithelial integrity^9,17,30,32^.

The transmembrane Crb3 protein has been reported to interact with Pals1/Patj and Par3/Par6/aPKC protein complex, contributing to establishment and maintenance of apical-basal polarity in epithelial cells. We observed that the absence of Crb3 in colonoids led to a notable reduction in Pals1, Patj and Par3 during junctional assembly. These findings suggest that in addition to its established role in controlling epithelial polarity, Crb3 also contributes to junction assembly^14,30,33^. The C-terminal cytoplasmic tail of Crb3 contains a FERM-binding domain (FBD) and a PDZ-binding motif (PBM). Previous studies identified that Crb3 associates with Pals1 and Par6 through the PBM, to establish apical-basal polarity and mature tight junctions in model epithelial cells^14,34^. Specifically, knockdown of Pals1 or Patj results in a leaky epithelial barrier associated with decreased junctional F-actin and discontinuous ZO-1 localization at TJs in epithelial cells^4,35^. We observed discontinuous ZO-1 labeling associated with disorganization of the perijunctional F-actin ring in primary murine IECs expressing the Crb3 mutant lacking PBM (Crb3ΔPDZ). Analogous to a Crb3a PBM mutant, putative mutations in the Crb3a mFBD (FBD mutant) failed to rescue uniform junctional localization of TJ protein (ZO-1) and organization of perijunctional F-actin ring. These findings suggest that both the FBD and PBM domains of Crb3a are required for the appropriate assembly and organization of the AJC and perijunctional F-actin ring.

Crb3 interacts with FERM domain proteins such as Ezrin, that link cell membrane proteins to the underlying actin cytoskeleton. Mice with global loss of Ezrin (Ezrin-/-) have been shown to display abnormal morphogenesis of intestinal villi and microvilli similar to mice with germline mutation of Crb3, suggesting that Crb3 controls intestinal morphogenesis during embryonic development via an Ezrin signaling pathway^36^. However, within intestinal epithelial cells, Ezrin localizes exclusively to the apical and ventral marginal zone (VMZ), that is spatially separated from Crb3 at AJC, suggesting that another ERM family member interacts with Crbs3 in the AJC. NF2, another member of the ERM family has been shown to localize in both TJ and AJ in MDCK cells^21^. This protein forms a complex with Pals1/Patj at TJs^21^ and directly interacts with α-catenin at AJ, thereby coupling the AJ to the underlying peri junctional F-actin ring in epithelial cells^20^. Using biochemical pulldown assays, we identified that Crb3 indeed interacts with the active form of NF2 via its FBD. In colonoids lacking Crb3, NF2 localization was no longer restricted to the AJC but instead spanned the lateral plasma membrane. This mislocalization suggests that the interaction between Crb3 and NF2 via FDB regulates NF2 localization at the AJC. Additionally, knockdown of NF2 in IECs resulted in a failure to establish barrier function, as evidenced by discontinuous localization of ZO-1 at TJs and disorganized perijunctional F-actin rings. These defects phenocopy Crb3 null and Crb3 FBD mutant expressing colonoids, highlighting the essential nature of Crb3-NF2 interaction during junctional assembly and establishment of the barrier.

Interestingly, our study found that while a Crb3 FBD mutant significantly decreased Crb3- NF2 interaction, the association with Pals1 was not altered. This observation aligns with previous findings that Crb3 associates with Pals1/Patj complex via the PBM domain. Despite the inability of the Crb3 FBD to interact with the Pals1/Patj complex, expression of this mutant did not rescue organization of the perijunctional F-actin ring and ZO-1 localization within TJs. Furthermore, it has been reported that Crb3 can associate with NF2 indirectly via Par3 through the PBM domain^20,37^. This dual mode of interaction of Crb3a with NF2; direct via FBD and indirect via PBM, could explain why expression of Crb3a mutant proteins lacking either FBD or PBM domains fail to restore junctional localization of ZO-1 and perijunctional F-actin. Together, these findings highlight the complex interactions of Crb3 with binding partners via FBD and PBM domains. Such interactions therefore facilitate the organization of the perijunctional actomyosin ring and, consequently, assembly of the AJC in intestinal epithelial cells.

Spatiotemporal controlled regulation of F-actin polymerization and myosin II-mediated contraction of the apical actomyosin ring is critical for AJC assembly and establishment of epithelial barrier function The small GTPase RhoA, in its active form, functions as a master regulator of organization and contractility of the apical actomyosin ring. Several studies have demonstrated that a balance of RhoA signaling is essential for the establishment and remodeling of the AJC and epithelial barrier integrity ^1,24,38,39^.

Our study revealed that the loss of Crb3 or NF2 in IECs increases F-actin and phosphorylated Myosin II (pMyosin II) density in the perijunctional F-actin ring. Inhibiting Myosin II hypercontractility with ROCK inhibitors (Y27632, H1152) or NMII ATPase inhibitor (Blebbistatin) in Crb3-null colonoids restored the perijunctional actomyosin ring organization and AJC protein localization. Live cell imaging showed higher RhoA activity at apical junctions in Crb3-null colonoids compared to controls. Additionally, Crb3-null cells exhibited higher junctional tension, indicated by vinculin localization at the AJC^28,29^. Our findings suggest that Crb3 loss enhances RhoA activity at apical junctions, leading to increased actomyosin contractility via ROCK-mediated Myosin II phosphorylation and greater junctional tension. F-actin polymerization changes may also result from actin-binding proteins like formins, activated downstream of RhoA. The mechanism linking Crb3 loss to increased RhoA activation at apical junctions is unclear but may involve altered activity of RhoGEF (p114RhoGEF) or RhoGAP (ARHGAP29), both associated with TJs and regulated by Crb3 partners Pals1 and Patj^11,40^. Further studies are needed to elucidate the pathways involved and to confirm the roles of these potential regulations in the context of Crb3.

The findings in this study highlight the critical role of Crb3 in regulating the spatiotemporal organization of apical junctions and the actomyosin cytoskeleton by modulating RhoA signaling, myosin II contractility, and junctional tension in epithelial cells during junction assembly. Proper organization of AJC and establishment of barrier function are essential for development during epithelial junction formation and in diseased states where epithelial junctions are remodeling following injury. Although studies on Crumbs3 and NF2’s involvement in inflammatory bowel disease (IBD) remain limited, our findings suggest that Crumbs3 and NF2 dysfunction may play a role in exacerbation of permeability defect during junction remodeling in inflammatory states such as IBD. A deeper understanding of these mechanisms may pave the way for novel strategies in managing and treating IBD.

## Methods and materials

### Antibodies and reagents

Used in this study are summarized in Supplemental table 1.

### Mice

*Crb^ERΔIEC^* and littermate control *Crb3^fl/fl^*mice were generated by breeding *Crb3^fl/fl^* ^9^ and Villin^ERT2-Cre^ in-house at the University of Michigan in Ann Arbor. 8-week-old *Crb3^ERΔIEC^* and control *Crb3^fl/fl^* mice were injected intraperitoneally with 1mg/100µL of tamoxifen (Sigma #T5648) for five consecutive days followed by a period of 30 days to rest. Mice were kept under specific pathogen-free (SPF) conditions with ad libitum access to normal chow and water. All experiments were approved and conducted following guidelines set by the University of Michigan Institutional Animal Care and Use Committee.

### Intestinal epithelial cells (IECs) and murine colonoids monolayer culture

SK CO-15 and primary murine colonoids were cultured as previous described^41^. Briefly, colon were harvested from *Crb3^ERΔIEC^* and *Crb3^fl/fl^* mice treated with tamoxifen and rested for 30days Harvested colons were subjected to chemical disruption using 50mM EDTA in 1x PBS (-) for 40minutes followed by mechanical shaking in shaking buffer (43.3 mM Sucrose and 54.9 mM Sorbitol in1x PBS(-)) to isolate colonic crypts. Colonic crypts were plated directly on culture plates coated with human collagen type IV (33µg/mL, Sigma #5533) and rh-Laminin (Thermo Fisher # A29248). Murine colonoids were maintained until the desired confluency in LWRN conditioned media supplemented with 50ng/mL human EGF (R&D Systems #236-EG) and antibiotics-antimycotic (Corning 30-003-Cl).

### Plasmids

siRNA targeting human NF2 and control (scrambled) siRNA were purchased from ThermoFisher (Silencer select siRNA Cat#4392421, Silencer select negative control siRNA Cat#4390844). Lentiviral plasmids Lentilox-RSV-Myc-Crb3a-CMV-GFP-puro-VSVG- Wt and mutants were made at the Vector Core of University of Michigan.

### siRNA transfection, exogenous Crb3 transfection, and lentiviral infections

Human model IEC, SKCO-15, and murine primary colonic epithelial cell monolayers were cultured either on Transwell permeable supports (Corning) or on tissue culture plastic chamber slides as described previously. Knockdown of Crb3 or NF2 in SKCO-15 cells was established by transfecting siRNA specifically targeting Crb3 or NF2 using Lipofectamine RNAiMAX (ThermoFisher #13778150). A scrambled and non-silencing siRNA was used to generate control cells. For ectopic expression of Myc-tagged Crb3, SKCO-15 cells were transfected with human pSEC Myc-*Crb3 wt and mutant constructs* using Lipofectamine 3000 (ThermoFisher #3000001); colonoids derived from *Crb3^ERΔIEC^* and *Crb3^fl/fl^* mice were infected with lentivirus expressing vector pLLV human Crb3a wt and mutants using 8ug/ml of polybrene (Millipore Sigma #TR-1003).

### Immunofluorescence

Human model IEC, SKCO-15, and murine colonoids were cultured on plastic chamber slides were fixed with 4% paraformaldehyde (PFA) and permeabilized with 0.2- 0.5% Triton X-100 for 10 minutes. Monolayers were blocked with 3% goat or donkey serum in PBS+ with 0.05% Tween-20 blocking buffer for 30 minutes. Primary antibodies were diluted in blocking buffer and cells were incubated overnight at 4°C. Cells were washed with PBS with 0.05% Tween 20 and fluorescently labeled secondary antibodies were diluted in blocking buffer followed by incubation for one hour at room temperature. Cells were washed and mounted in Prolong™ Gold anti-fade agent (ThermoFisher Scientific #P36930). Frozen colonic tissue sections (6-8µm) from *Crb3^ERΔIEC^*and *Crb3^fl/fl^* mice were fixed with 4% PFA, permeabilized with 0.5% Triton X-100 and immunolabeling performed as described above. High resolution fluorescent confocal images were captured using a Nikon A1 confocal microscope in the Microscopy & Image Analysis Laboratory Core at U-M.

### Immunoprecipitation and Western blotting

For immunoprecipitation studies, SKCO-15 naïve or SKCO-15 transfected with exogenous Myc-Crb3 wt and mutants were harvested in relaxation lysis buffer (10 mM HEPES, 10 mM NaCl, 3.5 mM MgCl2, 100 mM KCl, 1% Octyl-glucoside) supplemented with a cocktail of protease and phosphatase inhibitors. Epithelial lysates were centrifuged, and supernatants collected. Supernatants were pre-cleared with 50 µl of 50% protein A/G agarose beads (ThermoFisher Scientific, #20421) for 1 hour followed by incubating with 5 µg/ml antibodies or control IgG rotated overnight at 4° C. Immune complexes were precipitated by 25 µl of protein G Dynabeads (ThermoFisher Scientific, #1003D) for four hours. Immunoprecipitated complexes were washed before boiling in 2X NuPAGE LDS Sample Buffer (ThermoFisher Scientific #NP0007). Immunoprecipitates and input were then subjected to SDS-PAGE followed by immunoblotting.

### Permeability assay

Colonoids derived from *Crb3^ERΔIEC^* and *Crb3^fl/fl^*mice were grown on Transwell permeable supports ((0.4 μm pore-size, Corning, #3460) to confluence. TER-to-passive ion flow was measured continuously daily using an EVOMX voltmeter (World Precision Instruments, Sarasota, FL) for 6 days. For FITC dextran flux experiments, colonoids from *Crb3^ERΔIEC^*and *Crb3^fl/fl^* mice were grown on Transwell permeable supports till TER of *Crb3^fl/fl^* colonoids reached 600 Ω.cm^2^. After upper and lower Transwell compartments were washed twice with HBSS and pyruvate buffer (10 mM HEPES, pH 7.4, 1 mM sodium pyruvate, 10 mM glucose, 3 mM CaCl2, and 145 mM NaCl2) was placed in both of upper and lower compartments for 1 h at 37 °C. A freshly prepared solution of 4 kDa FITC-dextran dissolved in pyruvate buffer was added to the top chamber of the Transwells and incubated with gentle shaking for 2 h at 37 °C. Samples from the bottom chamber of the Transwells were collected as time points indicated, and the fluorescence intensity was measured with a fluorescent plate reader (BioTek, Winooski, VT).

### Lentiviral transduction and live imaging of active RhoA

For live cell imaging of active Rho, colonoids harvested from *Crb3^ERΔIEC^*and *Crb3^fl/fl^* mice were cultured on cell culture imaging dish with polymer coverslip bottom (ibidi, # 80136) coated with human collagen type IV (33µg/mL, Sigma #5533) and rh-Laminin (Thermo Fisher # A29248). Colonoids were infected with lentivirus packaged with vector, pLV-dimericTomato-2xrGBD (Addgene #176098) using 5-10ug/ml of polybrene (Millipore Sigma #TR-1003). Following 48hours post infection, colonoids were incubated with 200 nM Membright-488 dye (Millipore Sigma #SCT083) for 15minutes prior to imaging. High resolution live cell images of active RhoA and plasma membrane were captured using Zeiss LCM 980 Airyscan 2 equipped with a heated stage insert in the Microscopy & Image Analysis Laboratory Core at U-M. Z-stacks of 0.5µm thickness for a total of 5-8um were acquired to account for differences in the cell height and to acquire all cell-cell junctions in the field of view.

### Image analysis for active Rho

Active RhoA intensity in the colonoids derived from *Crb3^ERΔIEC^* and *Crb3^fl/fl^* mice were measured using ImageJ FIJI software. Using plasma membrane signal as a marker of cell-cell junctions, 25-pixel wide lines were drawn from vertex to vertex to cover the whole junction including bi-cellular and tri-cellular junctions. A similar line (25 pixel) was drawn in the cytoplasmic region close to each of the junctions to account for the cell-cell variable background signal. Relative fluorescence intensity of active Rho at the cell-cell junction was calculated by deducting the background signal from the junctional signal of the dTomato-2xrGBD for respective junctions.

### Statistical analysis

Averaged values are expressed as mean ± standard derivation (SD) or mean ± standard derivation (SEM). Statistical analysis was performed by two-tailed Student’s *t*- test or two-way ANOVA using GraphPad Prism software. *P* value less than 0.05 was considered significant. No samples were excluded from the analysis.

## Acknowledgements

The authors thank Dr. Roland Hilgarth for assistance in the generation of Crb3a WT and Crb3a mutant constructs, and lentivirus expressing Crb3a mutants and active Rho probe (dTomato- 2xrGBD). We also thank the Microscopy and Image Analysis Laboratory and the Unit for Laboratory Animal Medicine (ULAM) at the University of Michigan Medical School. This work was supported by the following grants: R01 DK129214 (AN and CAP) and DK59888 (AN).

---

**Supplemental Figure 1: Expression of Crb3 binding partners and histology of colonic mucosa.
(a)** Immunofluorescence labeling of colonic tissue sections of tamoxifen-treated *Crb3^fl/fl^* and *Crb3^ERΔIEC^* mice showing Crb3a, Pals1, Patj, Par3, and DAPI (nuclei/blue). Scale bar = 50 µm; **(b)** Hematoxylin and eosin (H&E) stained Swiss roll colonic tissue sections of tamoxifen- treated *Crb3^fl/fl^* and *Crb3^ERΔIEC^* mice showing comparable tissue architecture; Scale bar = 500 µm. Black box highlights an area that is enlarged in the bottom panel. Scale bar = 50 µm.

**Supplemental Figure 2: NF2 expression is independent of Crb3.** Immunoblotting of Crb3, NF2, pNF2S518 and β-actin (loading control) in colonoids derived from *Crb3^fl/fl^* and *Crb3^ERΔIEC^* mice.

**Supplemental Figure 3: Efficiency of NF2 knockdown in model IECs.** Immunofluorescence labeling of SK CO-15 IECs transfected with control scramble or two independent NF2 siRNA showing NF2 (green) and DAPI (nuclei/blue). Scale bar = 25 µm.

**Supplemental Figure 4: Inhibition of ROCK or pMLC activity rescues the general architecture of perijunctional F-actin ring and AJC.(a)** Immunofluorescence labeling of colonoids derived from *Crb3^ERΔIEC^* mice treated with vehicle (DMSO), ROCK inhibitor (H1152), and Non-Muscle Myosin II ATPase Inhibitor (Blebbistatin) showing p-MLC ^T^^18/S19^ staining; **(b)** Representative confocal images of perijunctional F-actin (Phalloidin), and AJC proteins (ZO-1 and E-Cadherin) in Crb3^fl/fl^ colonoids and Crb3^ERΔIEC^ colonoids treated with ROCK II inhibitor (Y27632, 50 µM) or vehicle (DMSO, 0.1%) for 30 minutes. Scale bar = 25 µm.

**Supplemental Figure 5:(a-b)** Representative confocal images of subconfluent colonoids derived from Crb3^fl/fl^ and Crb3^ERΔIEC^ mice immunostained for **(a)** Vinculin (rabbit anti-Vinculin, green) and ZO-1 (red); **(b)** Vinculin (mouse anti-Vinculin, green) and F-actin (red). White box represents zoomed in region shown on the right. Scale bar = 25 µm; **(c)** Representative image of sub- confluent Crb3^ERΔIEC^ colonoids expressing pLenti-dTomato-2xrGBD (pseudo colored as FIRE LUT) and stained with MemBright 488 plasma membrane dye (green), treated acutely with vehicle (water) and active Rho inhibitor (C3 transferase, 2µg/ml). Images shown are sum of Z-projections. Scale bar = 20 µm; **(Right)** Graph showing the relative intensity of active Rho at cell-cell junctions normalized to the background signal of every junction. Data are mean ± SEM. Each dot represents an independent junction from 2 independent experiments **** p <0.0001 by Mann-Whitney test. n=18 junctions for control (Crb3^ERΔIEC^); n=21 junctions for C3 transferase (C3T+Crb3^ERΔIEC^).

**Supplimentary table 1.**

| **Specificity** | **species** | **Suppliers** | **Catalog number** | **Dilution** |
| --- | --- | --- | --- | --- |
| Crb3 | rabbit | In-house | N/A | WB 1:4000; IF 1:500 |
| β-Actin | mouse | Sigma Aldrich | 5411 | WB 1:5000 |
| Merlin | rabbit | Cell Signaling Technology | 12888 | IF 1:100 |
| Merlin | rabbit | Cell Signaling Technology | 6995 | WB bad |
| Myc-tag | mouse | Cell Signaling Technology | 2276 | IP 1:100 |
| Myc-tag | rabbit | Cell Signaling Technology | 2278 | WB 1:2000 |
| Pals1 | rabbit | In-house | N/A | WB 1:1000; IF 1:200 |
| PARD3(Par3) | rabbit | Novus | NBP1-88864 | WB 1:1000; IF 1:100 |
| PATJ | rabbit | In-house | N/A | IF 1:100 |
| PATJ/INADL | rabbit | LifeSpan Bioscience | LS-C410011 | WB 1:1000 |
| p-MLCT18/S19 | rabbit | Cell Signaling Technology | 3674 | IF 1:250 |
| p-MLC S19 | rabbit | Cell Signaling Technology | 3671 | IF 1:250 |
| ZO-1 | mouse | ThermoFisher | 33-9100 | WB 1:1000; IF 1:250 |
| E-Cadherin | goat | R & D System | AF648 | WB 1:2000; IF 1:500 |
| Ezrin | mouse | ThermoFisher | 35-7300 | WB 1:1000 |
| β-Catenin | mouse | BD Bioscience | 610153 | WB 1:1000; IF 1:500 |
| Alexa Fluor™ 488 Phalloidin |  | ThermoFisher | A12379 | IF 1:400 |
| Alexa Fluor™ 555 Phalloidin |  | ThermoFisher | A34055 | IF 1:400 |
| Vinculin | rabbit | GeneTex | GTX109940 | IF 1:100 |
| Vinculin | mouse | ThermoFisher | 700062 | IF 1:100 |
| Calnexin | rabbit | Abcam | 22595 | WB 1:4000 |

