## Supplemental Figures 1-5 for "Crb3 and NF2: A dynamic duo that controls assembly of the apical junctions and barrier function via Rho/ROCK signaling"

### Supplemental figure 1

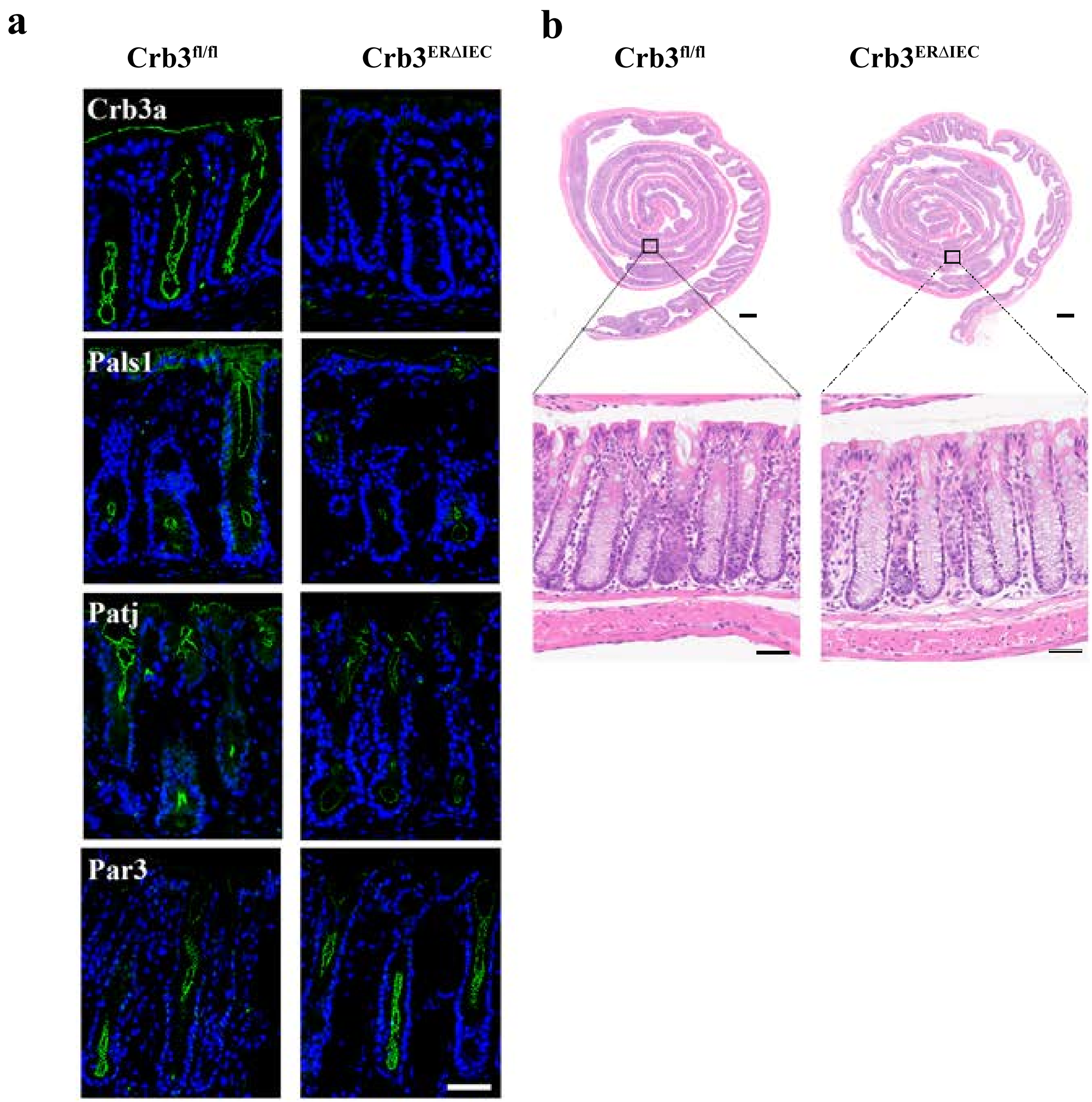

### Supplemental figure 2

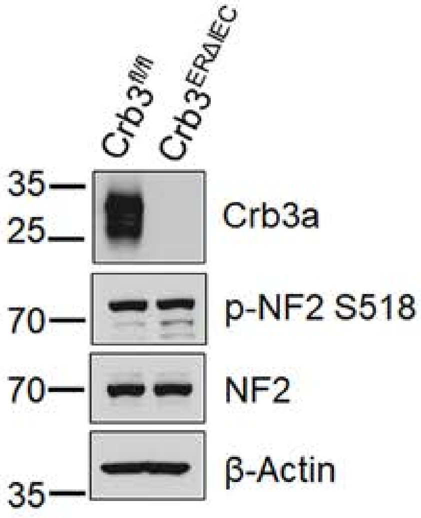

### Supplemental figure 3

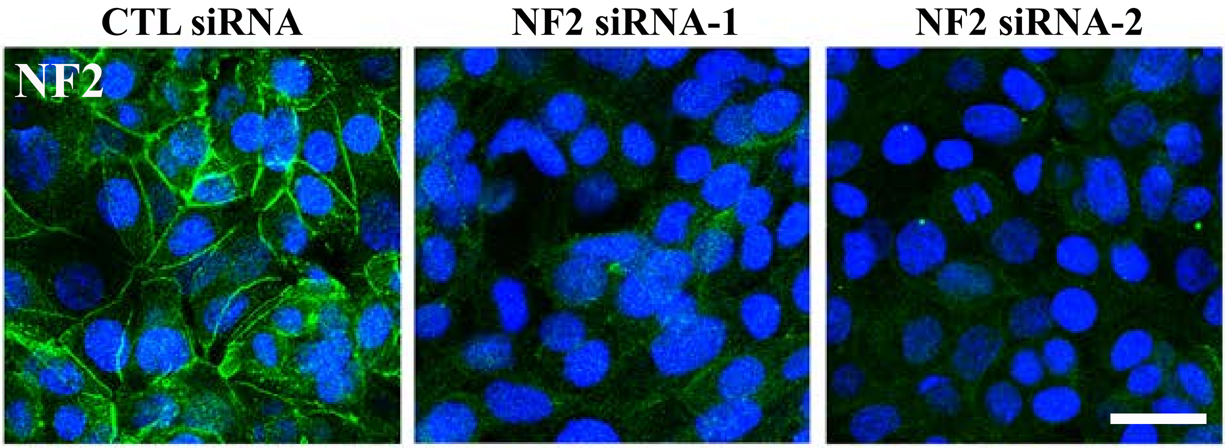

### Supplemental figure 4

**a**

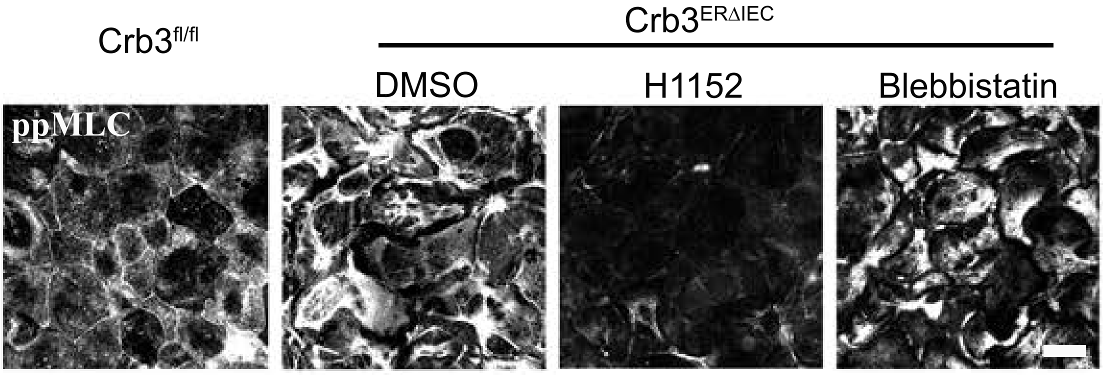


**b**

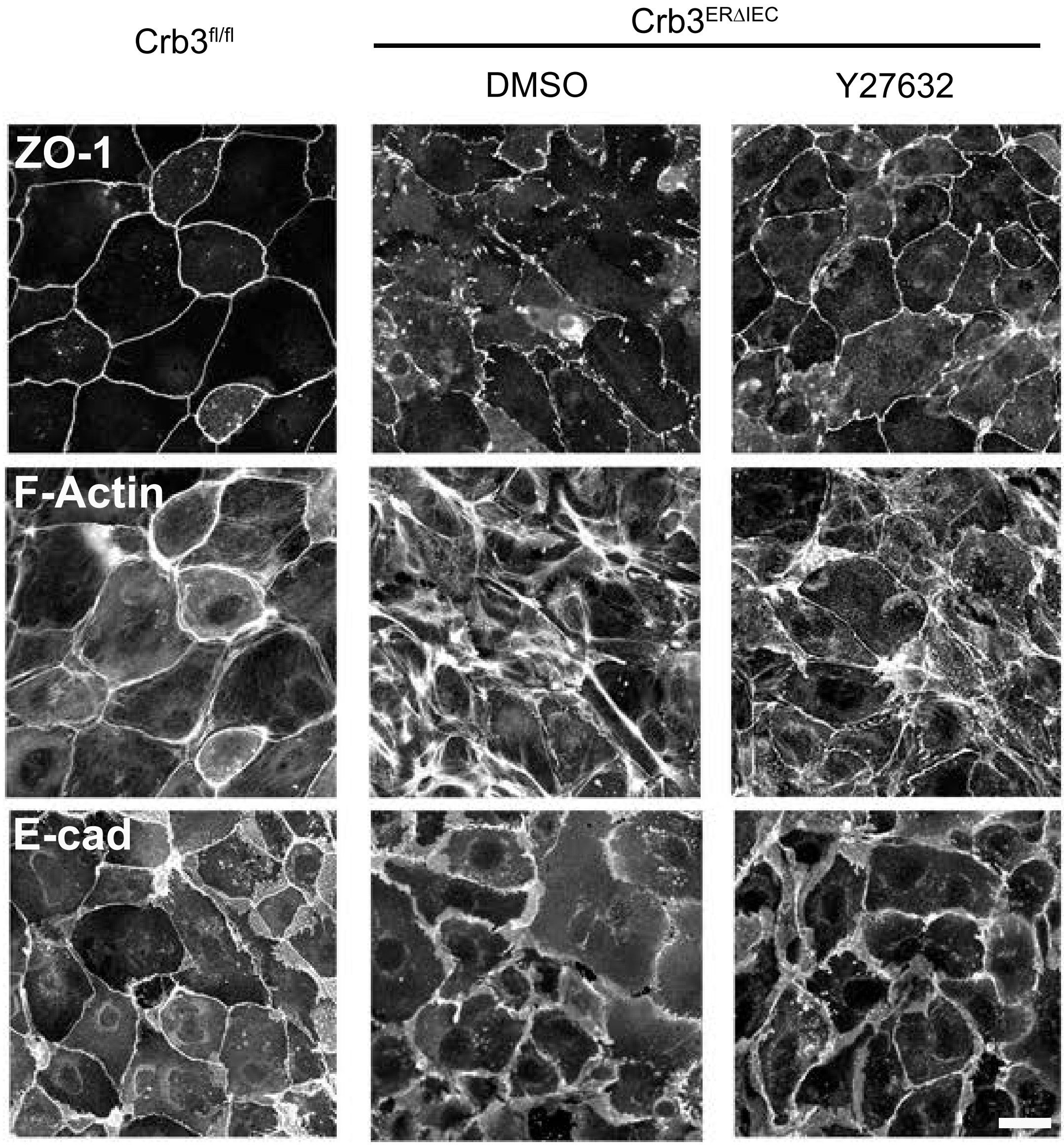

### Supplemental Figure 5

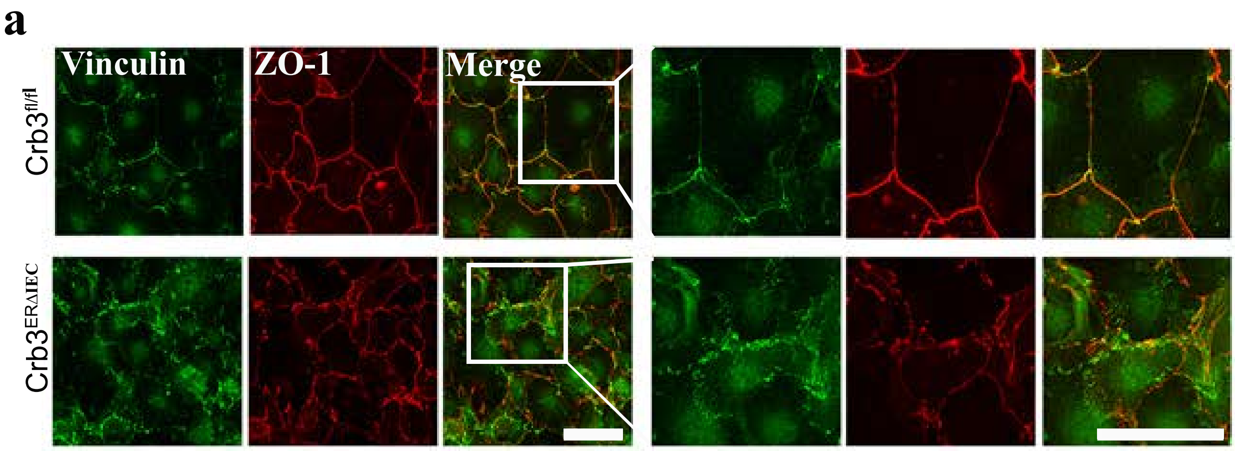


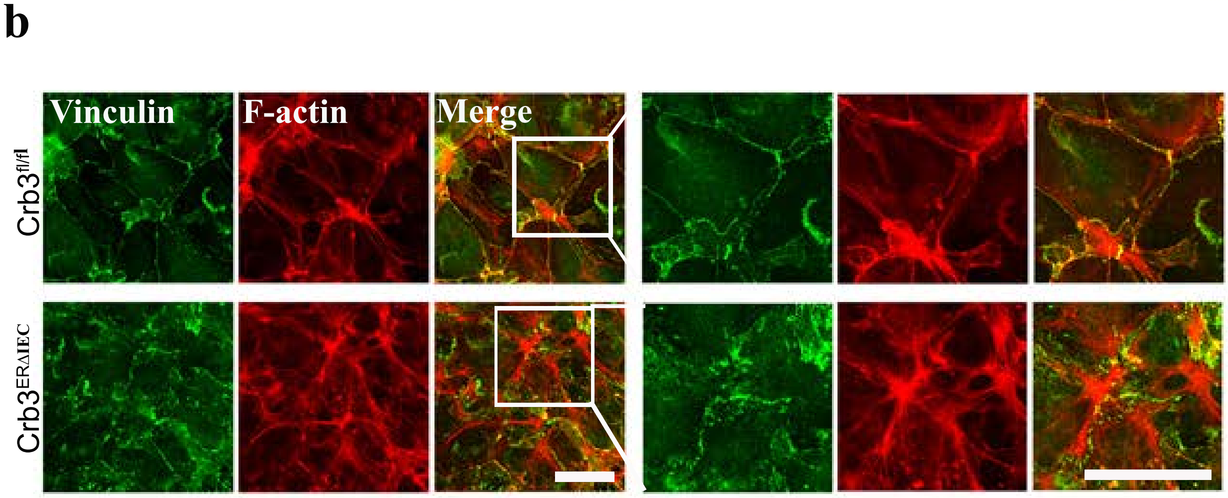


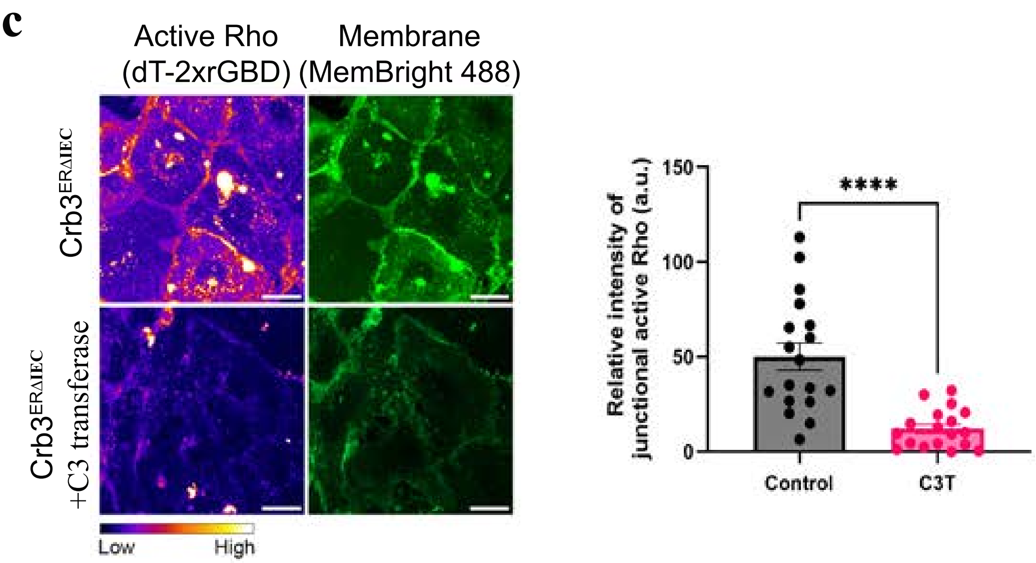
